# Paradoxic enhancement of mitochondrial capacity in aging-specific megakaryopoiesis from hematopoietic stem cells

**DOI:** 10.1101/2025.11.10.687744

**Authors:** Saran Chattopadhyaya, Stephanie Smith-Berdan, Sarah Huerta, E. Camilla Forsberg

## Abstract

Aging leads to quantitative and qualitative changes in platelet (Plt) production, with increased risk for thrombosis and other adverse cardiovascular events. Recent reports showed that aging promotes the emergence of non-canonical (nc) megakaryocyte progenitors (MkPs) directly from hematopoietic stem cells (HSCs), leading to the production of hyperactive Plts. The higher engraftment potential of ncMkPs compared to both young and old canonical (c)MkPs, contrasts with the functional decline of old HSCs. Emerging reports suggest that mitochondrial function critically regulates lineage commitment and cellular functionality, but how mitochondrial activity affects aging megakaryopoiesis is unknown. Here, we demonstrate that aged MkPs sustain unique mitochondrial activity, characterized by higher mitochondrial membrane potential, higher ATP content, and lower ROS levels compared to their younger counterparts. This contrasts with the dysfunctional mitochondrial state observed in old HSCs, suggesting lineage-specific organelle adaptations upon aging. Notably, we observed that the elevated mitochondrial capacity in aged MkPs is driven selectively by the age-specific ncMkPs. Paradoxically, in vivo pharmacological enhancement of mitochondrial activity in old mice reduced in situ Plt production, but increased Plt reconstitution by transplanted HSCs. These discoveries link uniquely regulated mitochondrial capacity to the intrinsic properties of age-specific MkPs, raising the possibility of therapeutic targeting to prevent aging-induced megakaryopoiesis.

**HIGHLIGHTS:**

- Aging-specific MkPs have elevated mitochondrial capacity, the inverse of aged HSCs
- Mitochondrial enhancement differentially alters platelet counts in young and old mice
- Enhancement of mitochondrial capacity increases platelet repopulation by both young and old HSCs

## INTRODUCTION

Aging is associated with profound and progressive alterations in the hematopoietic system, leading to increased susceptibility to a wide range of hematopoietic disease, including platelet (Plt) related disorders^1,2^. Hematopoietic stem cells (HSCs) are responsible for the constant supply of blood cells throughout life. Aging skews the hematopoietic system by altering the numbers and function of both HSCs and progenitors^3^. Old (o) or aged HSCs are characterized by reduced reconstitution capability^3^ and, as recently discovered by our lab, differentiation into an additional age-specific Plt production path^4^. This aging-progressive pathway operates in parallel with “canonical” Plt production that progresses from the HSCs through via multi- and oligopotent progenitor cell states throughout the lifespan. Using a double transgenic “*FlkSwitch*” lineage tracing mouse model^5^, our lab recently reported a noncanonical, age-progressive mechanism of Plt generation via a shortcut pathway directly from HSCs to megakaryocyte progenitors (MkPs). This additional “shortcut” pathway results in an age-specific expansion in the number of MkPs and Plts^4^. In contrast, no significant changes were observed in Plt production through the canonical (c)MkP pathway upon aging, which remains the primary source of Plt generation in young (y) adult mice ^4^. While HSCs functionally decline with age^3^, our studies revealed that age-specific, non-canonical (nc)MkPs produced directly from oHSCs generate hyperactive age-specific Plts and exhibit higher Plt repopulation capability upon transplantation. This increased functionality of ncMkPs upon aging sharply contrasts with the functional decline of oHSCs^3,4^. The mechanisms behind the superior capacity of ncMkPs are unknown, but essential to understand as they could provide novel strategies for reducing thrombotic risk.

Aging leads to various intrinsic cellular changes, including mitochondrial activity^6,7^. Emerging evidence suggests that mitochondria play a multifaceted role, encompassing energy production, cell fate decisions, and epigenetic regulation of stem cells^7,8^. Accumulating evidence indicates that yHSCs have high mitochondrial membrane potential (MMP) and lower levels of reactive oxygen species (ROS) compared to oHSCs^8–10^. Higher MMP and lower ROS levels equate to more robust stem cell function. Conversely, age-associated mitochondrial dysfunction leads to a decline in HSC functionality. Therapeutic interventions such as mitoquinol 10 (MitoQ) has been reported to rejuvenate aged HSCs by restoring mitochondrial activity^9^. While the effect of mitochondrial enhancement on megakaryopoiesis is unknown, separate studies have provided strong evidence for mitochondrial involvement in Plt function, including age-related changes and hyperreactivity^2,11–13^. How mitochondrial activity influence the two co-existing Plt production pathways in aging is still unclear. Thus, investigating the dynamics of mitochondrial function in oMkPs and oHSCs in comparison to younger counterparts offers critical insights into the mechanisms underlying healthy Plt production throughout life.

In the present study, we explore the possibility that changes in mitochondrial function may be the possible cause of the higher regenerative potential of oMkPs, especially age-specific ncMkPs. We investigated several mitochondrial parameters to understand the regulation of mitochondrial function across cell types and age. We also tested whether in vivo manipulation of mitochondrial capacity in vivo impacted quantity or quality of age-specific MkPs. These studies improve the understanding of age-related Plt production and suggest the possibility of developing future strategies to ensure balanced Plt production throughout life.

## RESULTS

### Aging induces megakaryocyte progenitor heterogeneity

Aging alters BM hematopoietic cellular frequencies and numbers and diminishes mitochondrial function^3,8^. To clarify the age-related changes in the frequency and number of HSCs and MkPs, we quantified cellular frequencies and numbers by flow cytometry. We observed a significant increase in HSC and MkP populations upon aging using defined phenotypic surface markers (**Fig 1A-B**). This is consistent with our own and others’ previous reports supporting that aging induces both HSC and MkP frequencies and numbers^3,14^. Our analysis also detected age-associated alterations within the MkP compartment. Our recent reports on MkPs showed that two distinct populations of MkPs exist in old mice; these can be differentiated by Tomato or GFP fluorescence in our FlkSwitch lineage tracing model^4,5^ or by differential surface expressions of CD48 and CD321: aging-independent, canonical cMkPs (CD48^+^CD321^-^) and age-progressive non-canonical MkPs (ncMkPs) (CD48^-^CD321^+^)^15^. Using the CD48/CD321 strategy, we did not observe a significant change in the frequency percentage in BM live cells and the number of cMkPs upon aging (**Fig 1C**), as expected^4,15^. In contrast, ncMkPs showed a significant increase in both frequency and numbers (**Fig 1C**). Altogether, these observations substantiated that the frequency and number of HSC and MkP populations increase upon aging, with the selective increase of ncMkPs, but not cMkPs, accounting for the expansion of the MkP pool upon aging. This observation is highly similar to our recent publication on the identification of age-specific megakaryopoiesis in the double transgenic FlkSwitch mouse model^4^, validating CD48 and CD321 as identifiers of cMkPs and ncMkPs in wild-type (WT) mice^15^. In summary, aging expands HSCs and MkPs, primarily through the selective increase of ncMkPs rather than cMkPs.

**Figure 1.**
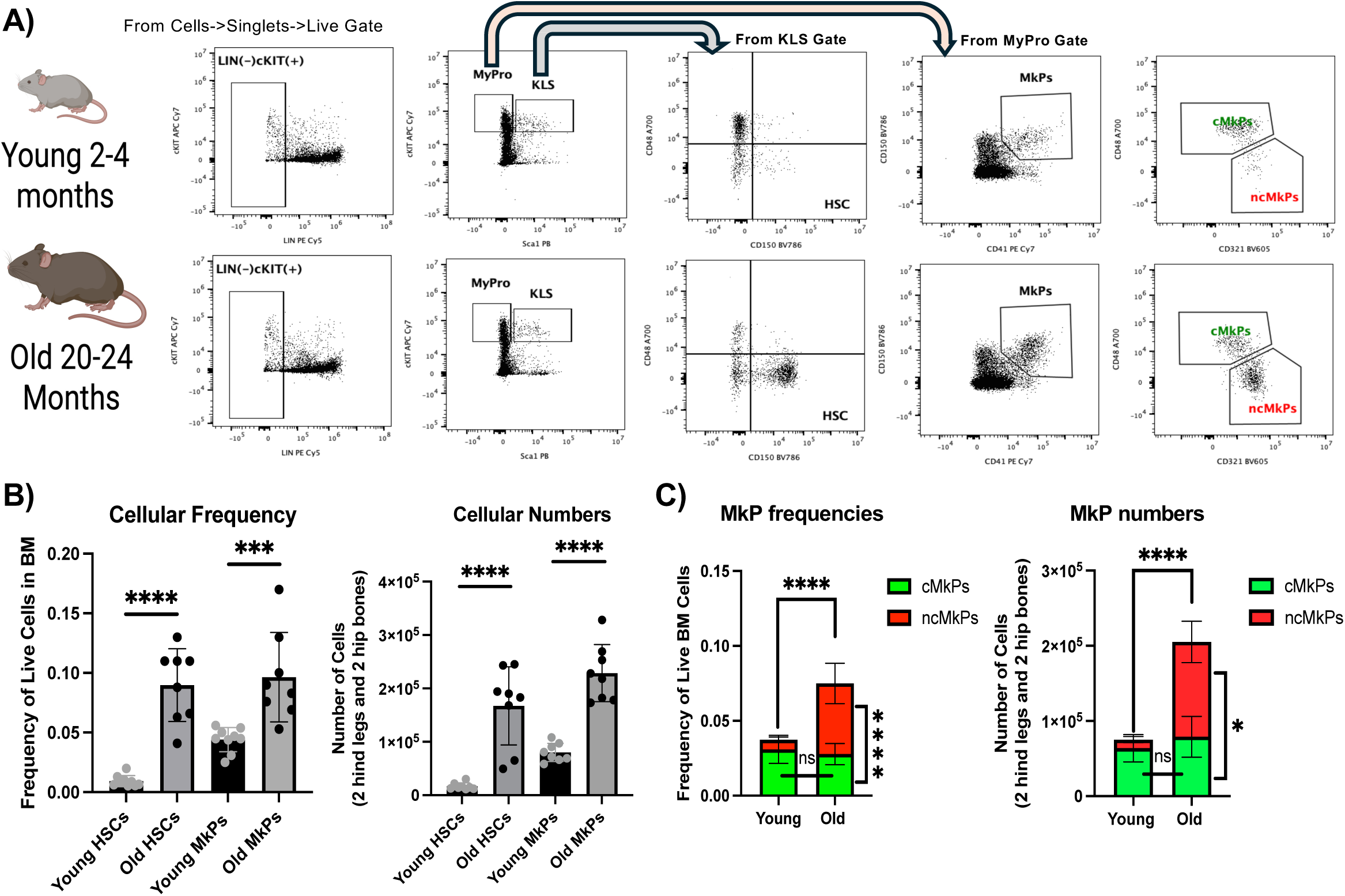
Hematopoietic stem cells, megakaryocyte progenitors and their subpopulations expand upon aging. **A.** Gating strategies for HSCs, MkPs, and canonical MkPs (cMkPs; young and old mice) and non-canonical MkPs (ncMkPs; old mice only). **B.** HSCs and MkPs increase upon aging. Frequency and total cell number analyses revealed significantly higher levels of HSCs and MkPs in the bone marrow (BM) of old mice compared with young mice. Student t-test. (n=8 in 5 independent experiments; ****P < 0.0001, ***P < 0.001) **C.** ncMkPs, but not cMkPs, increase upon aging Frequency and total cell number analyses revealed a significant increase in old ncMkPs upon aging, while there was no significant difference between young and old cMkPs. Student t-test. (n=6-7, ****P < 0.0001)

### Mitochondrial DNA and protein content change in HSCs and MkPs upon aging

Our recent publications reported that the aged ncMkPs have higher repopulation potential upon transplantation, which contrasts the sharp decline in HSC reconstitution of hematopoiesis upon aging^3,4^. Based on the reported links between HSC aging and mitochondrial capacity^9,16–18^, we hypothesized that mitochondria may also regulate the functionality of aged MkP populations. To test this, we assessed several key parameters of mitochondria in HSCs and MkPs and their subpopulations. Mitochondrial DNA content was measured as an indicator of mitochondrial biogenesis and abundance^19^. Mitochondrial mass was evaluated using MitoTracker Green (MTG) to capture changes in mitochondrial protein content^20^. Mitochondrial membrane potential (MMP) was quantified with tetramethylrhodamine ethyl ester (TMRE) as a measure of mitochondrial integrity and energy-coupling efficiency^21^. Cellular bioenergetic capacity was assessed through ATP quantification, while oxidative stress was examined via mitochondrial ROS production (MitoSOX) and global cellular ROS (DCFDA)^9^. Together, these parameters provide a multidimensional view of how mitochondrial function is altered with age in HSCs and MkP subpopulations.

First, to understand how aging affects mitochondrial numbers in HSCs and MkPs and their subpopulations, we quantified the mitochondrial DNA content, a measure of the number of mitochondria inside the cell^19^. Using quantitiative mitochondrial-specific PCR, we showed that both oHSCs and oMkPs have higher mitochondrial DNA (mtDNA) content than their young counterparts. Interestingly, there was no significant difference between young and old cMkPs while old ncMkPs showed a significantly higher mitochondrial to nuclear DNA ratio compared to old cMkPs (**Fig 2A**). These observations indicate that oHSCs and oMkPs possess greater mitochondrial abundance than their young counterparts, which in MkPs is driven primarily by higher levels in old ncMkPs compared to old cMkPs.

**Figure 2.**
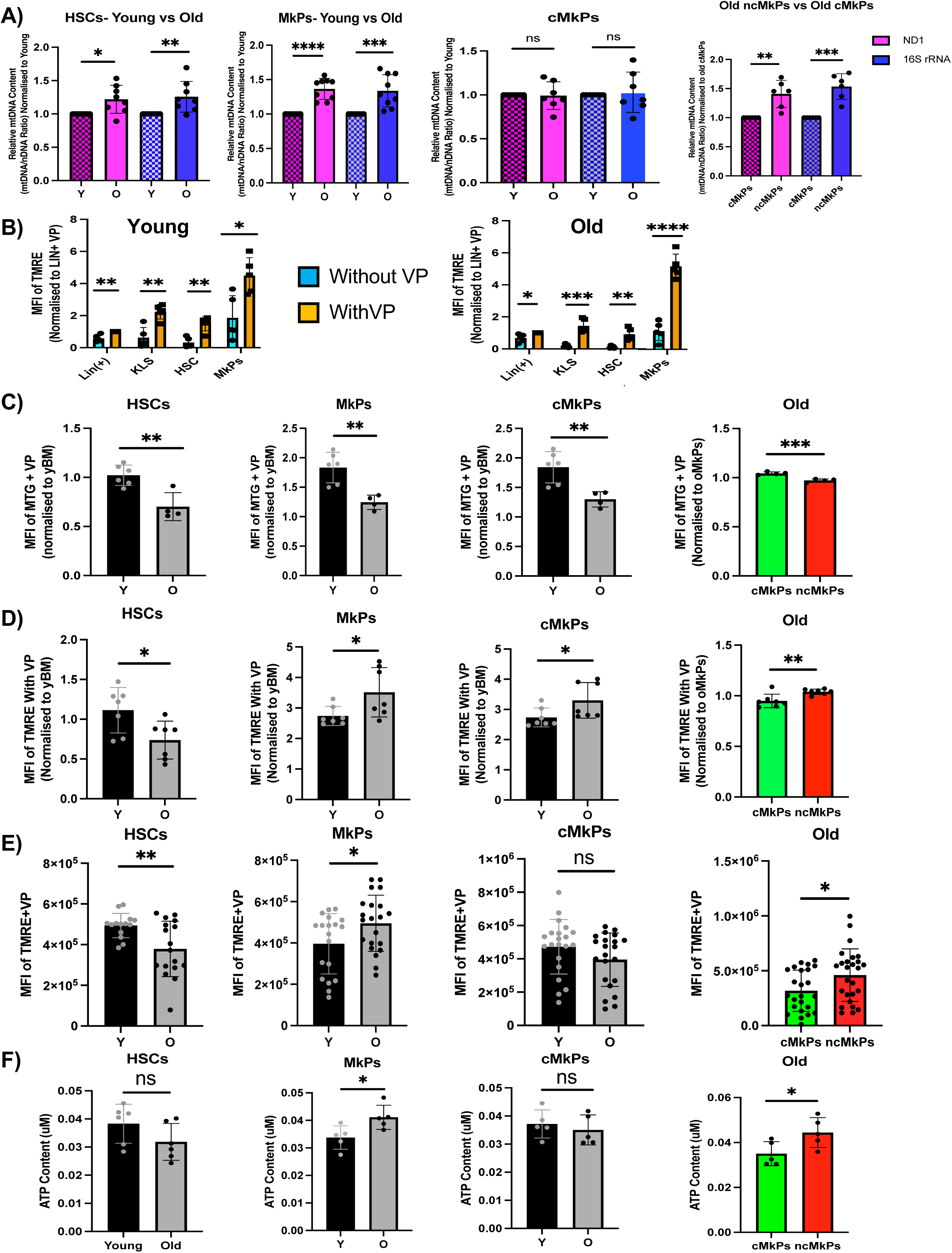
Aging induces opposing changes in mitochondrial activity in HSCs and MkPs, primarily driven by ncMkPs. **A.** oHSCs and ncMkPs have higher mitochondrial DNA content compared to young or co-existing counterparts. Relative qPCR quantification of mitochondrial DNA (mtDNA) was performed on samples from young (2-4 months; pattern fill) and old (18-24 months; solid/no pattern) mice. The ND1 and 16S genes, representing stable regions of mtDNA, were amplified and normalized to the nuclear hexokinase (HK) gene. A decrease in the mtDNA/nDNA ratio in young populations indicates a reduction in mtDNA copy number. Student’s t-test (n= 6-7 in 5 independent experiments, ****P < 0.0001, ***P < 0.001, **P < 0.01, *P < 0.05). **B.** Verapamil increases dye retention in young and old HSCs and MkPs. Effect of Verapamil on TMRE staining. MFI of MitoSox in absence of VP (cyan) normalised to MFI in presence of VP (orange) in BM. Student’s t-test. (n = 5 in 3 independent experiments; ****P < 0.0001, ***P < 0.001, **P < 0.01, *P < 0.05). **C.** Mitochondrial mass decreases upon HSC and MkP aging. Analysis of age-associated changes in mitochondrial mass measured by MitoTracker Green. Both young HSCs and MkPs showed significantly higher MFI of MTG. Plotted her as bar diagram. Student’s t-test (n= 4-6 in 3 independent experiments, ***P < 0.001, **P < 0.01). **D.** Mitochondrial membrane potential decreases in HSCs but increases in MkPs upon aging. Analysis of age-associated changes in mitochondrial membrane potential measured by TMRE. HSCs and MkPs showed opposite phenomena. Plotted as bar diagram. Student’s t-test (n= 7 in 5-6 independent experiments, **P < 0.01, *P < 0.05). **E.** Mitochondrial membrane potential is lower in cultured oHSCs compared to yHSCs, but higher in cultured oMkPs and ncMkPs. In-vitro comparison of MMP between young and old HSCs, young and old MkPs, young and old cMkPs and old cMkPs and old ncMkPs. HSCs were culture for 3 days and MkPs were cultured for 7 days. Student’s t-test. (n = 3 independent experiments; ****P < 0.0001, ***P < 0.001, **P < 0.01, *P < 0.05). **F.** ATP content is higher in age-specific ncMkPs compared to young and old cMkPs. Analysis of age-associated changes in cellular ATP content after sorting 1500 cells. Although there is no significant difference between young HSCs and old HSCs, however, old MkPs showed significantly higher ATP content compared to young MkPs. Significantly, old ncMkPs showed higher ATP content compared to old cMkPs. No significant difference between young and old cMkPs. Student’s t-test (n= 5-6 in 5 independent experiments, **P < 0.01, *P < 0.05).

We then evaluated dye-based mitochondrial activity in young and old HSCs and MkPs and their subpopulations. Because the presence of xenobiotic efflux pumps often leads to the inaccurate measurement of mitochondrial activity as cells can expel mitochondrial staining dyes^22^, we compared the effect of the efflux inhibitor verapamil (VP) on dye retention ability. Consistent with previous reports^9^, we observed that treatment with VP led to a significant increase in the median fluorescence intensity of mitochondrial tracking dyes across several distinct cell types, compared to Lin⁺ populations without VP treatment, irrespective of age (**Fig 2B**). Like HSCs^9^, MkPs also showed significant increases in MFI (median fluorescence intensity) for TMRE intensity with VP compared to the no VP-treated MkPs population (**Fig 2B**). Thus, to accurately measure efflux-independent mitochondrial activity, we subsequently used VP in our dye-based mitochondrial activity detection experiments (MTG, TMRE, and MitoSox).

Interestingly, quantification using MTG, a thiol-reactive, mitochondrial protein-binding fluorescent dye that represents mitochondrial mass^23^, showed that both yHSCs and yMkPs have higher MFI than their older counterparts (**Fig 2C**). This age-related decrease was also observed for both GFP+ and Tom+ oMkP subpopulations. While seemingly at odds with higher mtDNA content upon aging, this pattern is consistent with previous reports for HSCs ^22,24^, indicating that mitochondrial DNA and protein content are not directly correlated.

### Aging leads to inverse changes in mitochondrial capacity in HSCs and MkPs

Next, to better understand mitochondrial function upon aging, we compared MMP between the young and old HSCs and MkPs and their subpopulations by using TMRE dye. TMRE is a cell-permeant, cationic fluorescent dye that reversibly accumulates inside the mitochondria of living cells due to the negative charge of the mitochondria compared to the cytosol ^21^. Thus, the loss of the dye from mitochondria or a decrease in fluorescence intensity represents lower MMP, which ultimately directs lower energy (ATP) production. In accordance with previously published data^9^, we observed that yHSCs showed higher MMP compared to oHSCs (**Fig 2D**), consistent with the greater cellular functionality of yHSCs ^3,9^. Surprisingly, when we compared the MMP between y and o MkPs, the opposite outcomes were observed: oMkPs showed higher MMP than yMkPs (**Fig 2D**). Furthermore, old ncMkPs showed a significant increase in MMP compared to old cMkPs (**Fig 2D**), while old cMkPs also showed significantly higher MMP than young cMkPs (**Fig 2D**). To further investigate whether variations in mitochondrial function affect in vitro cellular proliferation, we sorted and cultured both young and old HSCs and MkPs, including their subpopulations. Upon in-vitro culture (3 days for MkPs and 7 days for HSCs), we observed that yHSCs and oMkPs have higher MMP compared to oHSCs and yMkPs, respectively (**Fig 2E**). Additionally, old ncMkPs showed higher MMP than old cMkPs, while young and old cMkPs showed no significant difference in the MFI of the membrane potential dye (**Fig 2E**). These results suggest that mitochondrial energetics are directly linked to proliferative output and provide additional support for our previous observation that old MkPs, especially old ncMkPs, maintain enhanced expansion capacity in vitro compared to their canonical counterparts, irrespective of age.

As MMP ultimately leads to cellular energy production (ATP), we measured ATP content directly. Consistent with prior data^25^, oHSCs showed no significant difference compared to yHSCs (**Fig 2F**). In contrast, MkPs showed robust age-related changes. oMkPs had higher ATP content than their younger population (**Fig 2F**). Importantly, while there was no significant difference between young and old cMkPs (**Fig 2F**), old ncMkPs displayed significantly higher ATP content than old cMkPs (**Fig 2F**) This divergence in aging-related ATP content between HSCs and MkPs highlights cell type-specific adaptations of mitochondrial activity in aging and provides new insight into how progenitor cells, but not HSCs, undergo mitochondrial activity reshaping with age. Combined with the increase in MMP, the greater ATP content of aged MkPs demonstrate that the decline in mitochondrial functionality with aging is not a universal phenomenon.

### In contrast to old HSCs, age-specific MkPs have reduced levels of ROS

Next, we measured reactive oxygen species (ROS) levels to determine whether changes in MMP and ATP content were associated with an altered redox status. Importantly, lower ROS levels are known to support higher mitochondrial function as well as cellular activity^26,27^. To ensure that we can accurately measure ROS levels using MitoSox, a dye that measures mitochondrial superoxide (ROS)^28^, we first used TBHP (Tert-butyl Hydroperoxide, a ROS enhancing agent^29^). TBHP-treated samples showed the expected increase in MitoSox values (**Fig 3A**), demonstrating the ability of this assay to quantify relative ROS levels. When applying the MitoSox assay to HSCs isolated from young and old mice, we observed a trend toward elevated ROS levels in oHSCs relative to yHSCs (**Fig 3B**). This observation supports earlier studies^8,30–32^ reporting that elevated oxidative stress is a defining feature of oHSCs and represents a major intrinsic factor contributing to their functional decline and loss of regenerative capacity. In contrast to oHSCs, oMkPs showed significantly lower ROS values than yMkPs (**Fig 3B**). Excitingly, this difference was primarily driven by ncMkPs, as young and old cMkPs did not exhibit differential ROS levels, whereas old ncMkPs showed significantly lower ROS values than old cMkPs (**Fig 3B**). To further characterize mitochondrial ROS during cellular proliferation, we measured MitoSox levels after sorting and culturing both young and old HSCs and MkPs, including their subpopulations. The patterns were consistent with the freshly isolated cells: oHSCs and yMkPs have higher ROS compared to yHSCs and oMkPs, respectively (**Fig 3C**). Additionally, old ncMkPs showed lower ROS than old cMkPs, while young and old cMkPs had similar levels of MitoSox (**Fig 3C**).

**Figure 3.**
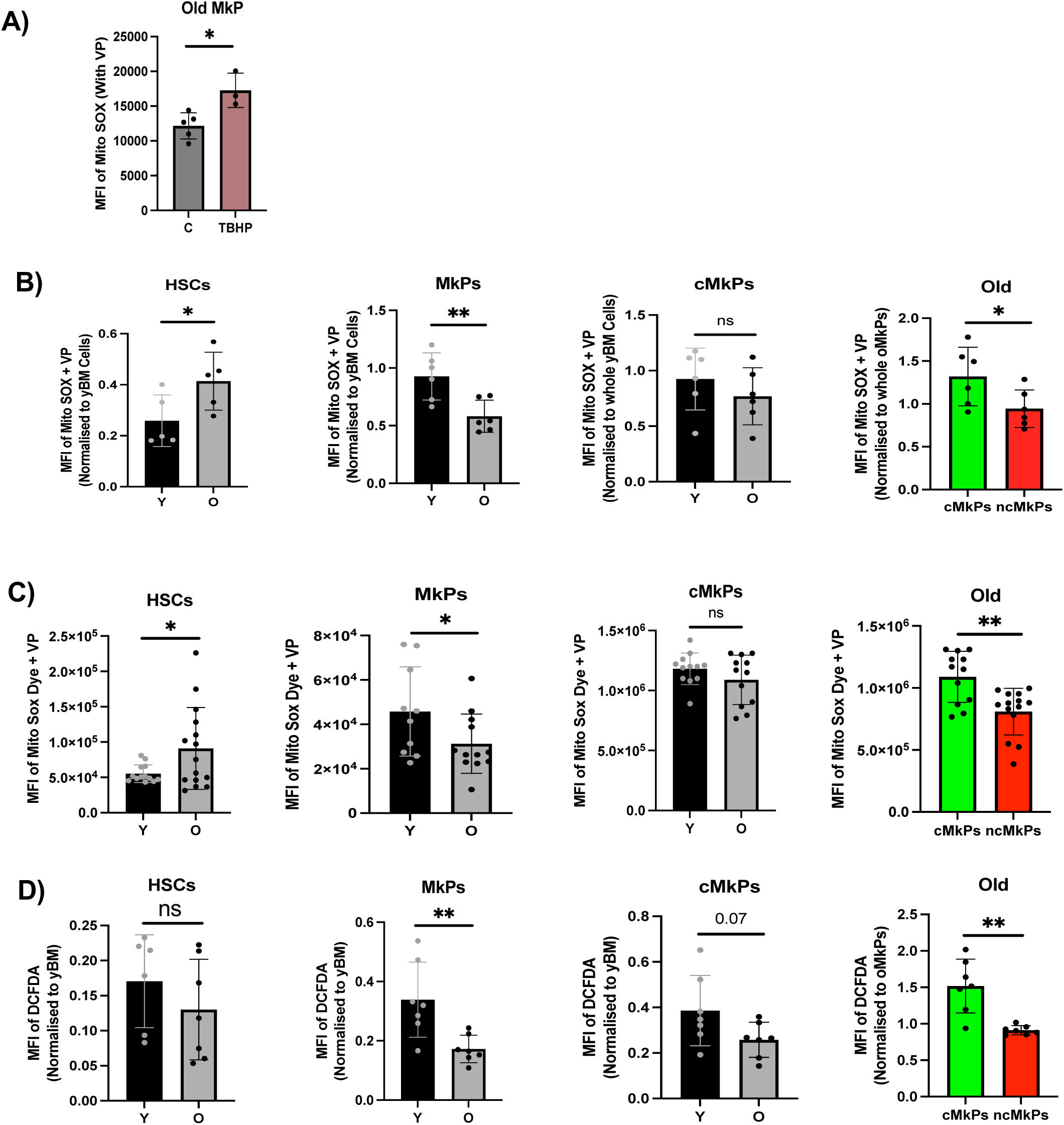
Aging leads to opposing changes in redox state of HSCs and MkPs. **A.** The ROS-enhancing reagent TBHP induces measurable alterations in MitoSox levels. Treatment with TBHP led to the expected increase in MitoSox levels. Comparison of untreated and positive control (TBHP) for MitoSox analysis. Student’s t-test (n= 3-4 in 3 independent experiments, *P < 0.05). **B.** Aged HSCs accumulate mitochondrial ROS, while aged MkPs have lower ROS levels due to ncMkPs. Analysis of age-associated changes in mitochondrial ROS (measured by Mito Sox). HSCs and MkPs showed opposite phenomena. Plotted as bar diagram. No significant difference between young and old cMkPs, while old ncMkPs showed significantly lower values compared to old cMKPs. Student’s t-test (n= 5-6 in 5 independent experiments, **P < 0.01, *P < 0.05). **C.** Aged HSCs cultured in vitro have higher ROS levels than yHSCs, while cultured ncMkP lowers ROS levels compared to young and old cMkPs.In-vitro comparison of MitoSox between young and old HSCs, young and old MkPs, young and old cMkPs and old cMkPs and old ncMkPs. Upon culture HSC and MkPs are holding up their ROS difference between young and old. Student’s t-test. (n = 3 independent experiments; ****P < 0.0001, ***P < 0.001, **P < 0.01, *P < 0.05). **D.** Age-specific ncMkPs are responsible for lower ROS levels upon MkP aging. Analysis of age-associated changes in cellular ROS (measured by DCFDA). Although no significant differences were observed between young and old HSCs, old MkPs showed lower ROS content than young MkPs. Plotted as bar diagram. No significant difference between young and old cMkPs, while old ncMkPs showed significantly lower values compared to old cMKPs. Student’s t-test (n= 6-8 in 5 independent experiments, **P < 0.01, *P < 0.05).

Similar observations were made when we compared overall intracellular ROS levels using DCFDA, which generates a green fluorescent signal after being oxidized by ROS^9^. While we did not observe a difference in cellular ROS between young and old HSCs, we found that oMkPs showed significantly lower intracellular ROS than yMkPs (**Fig 3D**). As with mitochondrial ROS, this difference was driven primarily by old ncMkPs (**Fig 3D**). Together, these data suggest that age-related increases in ROS are a defining feature of HSCs dysfunction, whereas old MkPs, and in particular ncMkPs, appear to resist this oxidative burden. This underscores a fundamental difference in redox homeostasis between these progenitor populations upon aging.

In summary, our findings indicate that aging does not result in a uniform decline in mitochondrial function across hematopoietic populations. Instead, mitochondrial regulation diverges sharply between HSCs and MkPs, with old ncMkPs uniquely sustaining and even enhancing mitochondrial capacity upon aging, positioning ncMkPs as a distinct, age-adapted progenitor population.

### Manipulation of in vivo mitochondrial activity differentially alters Plt counts in young and old mice

The inverse trajectories of mitochondrial potential of HSCs and MkPs upon aging raise the possibility that the higher MMP, higher ATP content, and lower ROS are responsible for their contrasting decline (aged HSCs) and enhancement (ncMkPs) in reconstitution potential upon aging^4,15^. Notably, the age-associated changes seen in MkPs are mainly attributable to the altered dynamics of the ncMkPs population. Evidence from others suggest that manipulating mitochondrial activity, including those affecting MMP, can influence HSC fate decisions and regenerative capacity^9,18^. Therefore, enhanced MMP is not only a readout of mitochondrial activity but also a functional regulator of stem cell functions. To evaluate whether short-term in vivo manipulation of mitochondrial activity, especially MMP, changes the production of mature blood cells in the PB, we next manipulated MMP levels in vivo. Mitoquinol Q (MitoQ), a mitochondrial-targeted coenzyme-Q10, has been reported to increase MMP and decrease ROS^33^. Notably, the effect of MitoQ on megakaryopoiesis is not known. Given that no surface markers have been identified to distinguish the two Plt pools upon aging in the wildtype mouse, we used FlkSwitch mice to understand the effect of mitochondrial functional enhancement on the three Plt populations that can be distinguished in this model (GFP+ yPlts, GFP+ oPlts and Tom+ oPlts). We treated young and old FlkSwitch mice with MitoQ for 5 days (**Fig 4A**). We observed that successive MitoQ injections significantly decreased the total body weight of both young and old mice (**Fig 4B**). Although our peripheral blood (PB) data analysis revealed no change in Plt numbers in the sham-treated groups of young and old individuals (data not shown), we observed a significant increase in Plt numbers in young mice upon 5 days of MitoQ treatment (**Fig 4C**). Surprisingly, we observed an opposite effect of mitochondrial functional enhancement in old mice: a significant decrease in Plt numbers (**Fig 4C**). When we further fractionated the two coexisting Plt populations in old mice, we found that although the number of GFP+ oPlts did not change upon 5 days of MitoQ treatment, there was a significant decrease in Tom+ oPlts (**Fig 4D**). Analysing the changes of GFP and Tom Plt percentages, we found that even though the proportions of Tom:GFP Plts varied among individuals, every old MitoQ-treated animal showed a reduction in Tom+ output of oPlts between the start and end of treatment (**Fig 4E**. This was accompanied by an increased proportion of GFP+ oPlts (**Fig 4E**). Combined with the reduced total numbers of Tom+ oPlts (**Fig 4D**), these data suggest that MitoQ treatment may selectively reduce the production of age-specific Plts from oHSCs. To test whether this may be due to a more general decline in HSC mature cell production upon MitoQ treatment, we quantified additional cell populations in PB. In the same cohorts of MitoQ-treated mice, but not in the sham-treated groups, the number of erythroid cells increased (**Fig 4F**) and B cells decreased (**Fig 4G**) in both young and old mice, with little to no changes in myeloid and T cell numbers (**Fig 4H-I**). Together, our data demonstrate that short-term MitoQ treatment exerts divergent effects on Plt populations in young versus old mice, with opposite outcomes depending on age. In contrast, the response of immune cells, such as B cells, remained largely consistent across age groups, suggesting that mitochondrial-targeted interventions may differentially influence existing mature hematopoietic lineages while preserving certain immune cell dynamics. Our data suggest that short-term pharmacological manipulation of mitochondrial function alters peripheral blood composition in an age-dependent manner. These findings indicate that mitochondrial activity influences lineage differentiation and progenitor behavior differently in young and old hematopoietic systems, highlighting age-related changes in mitochondrial responsiveness.

**Figure 4.**
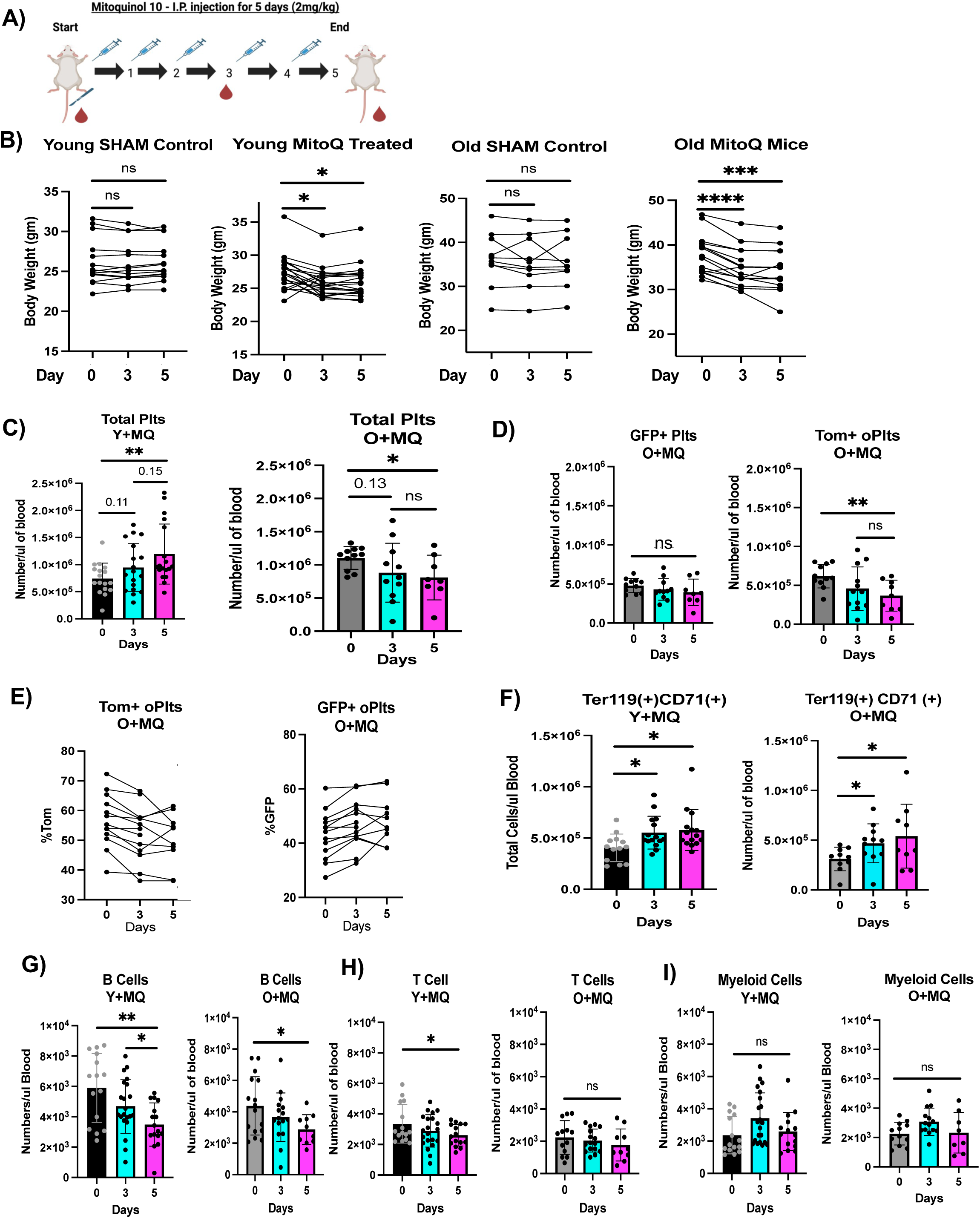
Enhancement of Mitochondrial Activity leads to divergent effects on Plt numbers in young and old mice. **A.** Schematic representation of 5-day in vivo MitoQ (MQ) treatment. **B.** Both young and old mice lose weight upon MitoQ treatment. Comparison of total body weight during 5 days of MQ treatment. One-way ANOVA mixed model – Tukey correction. (n = 8-14 in 3 independent experiments; ****P < 0.0001, ***P < 0.001, **P < 0.01, *P < 0.05). **C.** Plt counts increase in young mice but decrease in old mice upon MitoQ treatment. Number of platelets in PB before MQ treatment vs 3 and 5 days of MQ treatment. Young before-treatment (black) and old before-treatment (grey) treated mice at the beginning of treatment, compared to Y+MQ and O+MQ mice at the end of a 5-day treatment (purple), showed significantly increased and decreased numbers, respectively. Student’s t-test. (n = 11-16 in 3 independent experiments; **P < 0.01, *P < 0.05). **D.** Tom+ oPlts, but not GFP+ oPlts, decreased in response to MitoQ. Changes in the number of GFP+ and Tom+ Plt in PB of before treatment vs 3 or 5 days of MQ treatment in the same old mice as in (**C**). Student’s t-test. (n = 11-16 in 3 independent experiments; **P < 0.01, *P < 0.05). **E.** Percentage of GFP+ and Tom+ Plt changes in PB of individually measured old mice before (0), in the middle (3), and at the end of treatment (5). Student’s paired t-test. (n = 13). **F-I.** Alterations in erythroid, myeloid, B and T cells upon MitoQ treatment. Changes in the numbers of Ter119(+)CD71(+) erythroid cells (**F**), B cells (**G**), T cells (**H**), and myeloid cells (**I**) in the PB of before treatment vs 3 and 5 days of MitoQ treatment in both young and old mice. Student’s t-test. (n = 11-16 in 3 independent experiments; **P < 0.01, *P < 0.05).

### Mitochondrial potentiation alters HSC and MkP numbers and reconstitution potential

Our results showed that manipulation of mitochondrial function can lead to significant alterations in the number of specific PB cell populations. Such changes often arise due to changes in stem and progenitor cells in the BM. We therefore measured the numbers of HSCs and MkPs after 5 days of MitoQ treatment. Because Plt numbers increased in young MitoQ-treated mice, we anticipated an increase in HSCs and/or MkPs in young MitoQ-treated mice. Surprisingly, neither HSC nor MkP pools changed (**Fig 5A**). Additionally, while Plt numbers, in particular Tom+ Plts, decreased in old mice, we did not observe a significant difference in Tom+ oMkPs compared to the sham-treated group (**Fig 5B**). Interestingly, both oHSCs and GFP+ oMkPs increased in numbers (**Fig 5B**). The increase in oHSC numbers was particularly surprising, because we had hypothesized that MitoQ-mediated rejuvenation of oHSCs might reduce the oHSC pool to approach the smaller numbers of HSCs present in young mice. Collectively, these data suggest that rather than affecting Plt counts via influencing HSC and MkP numbers, the primary effects of MitoQ may be to alter HSC and MkP functional capacity via metabolic rewiring.

**Figure 5.**
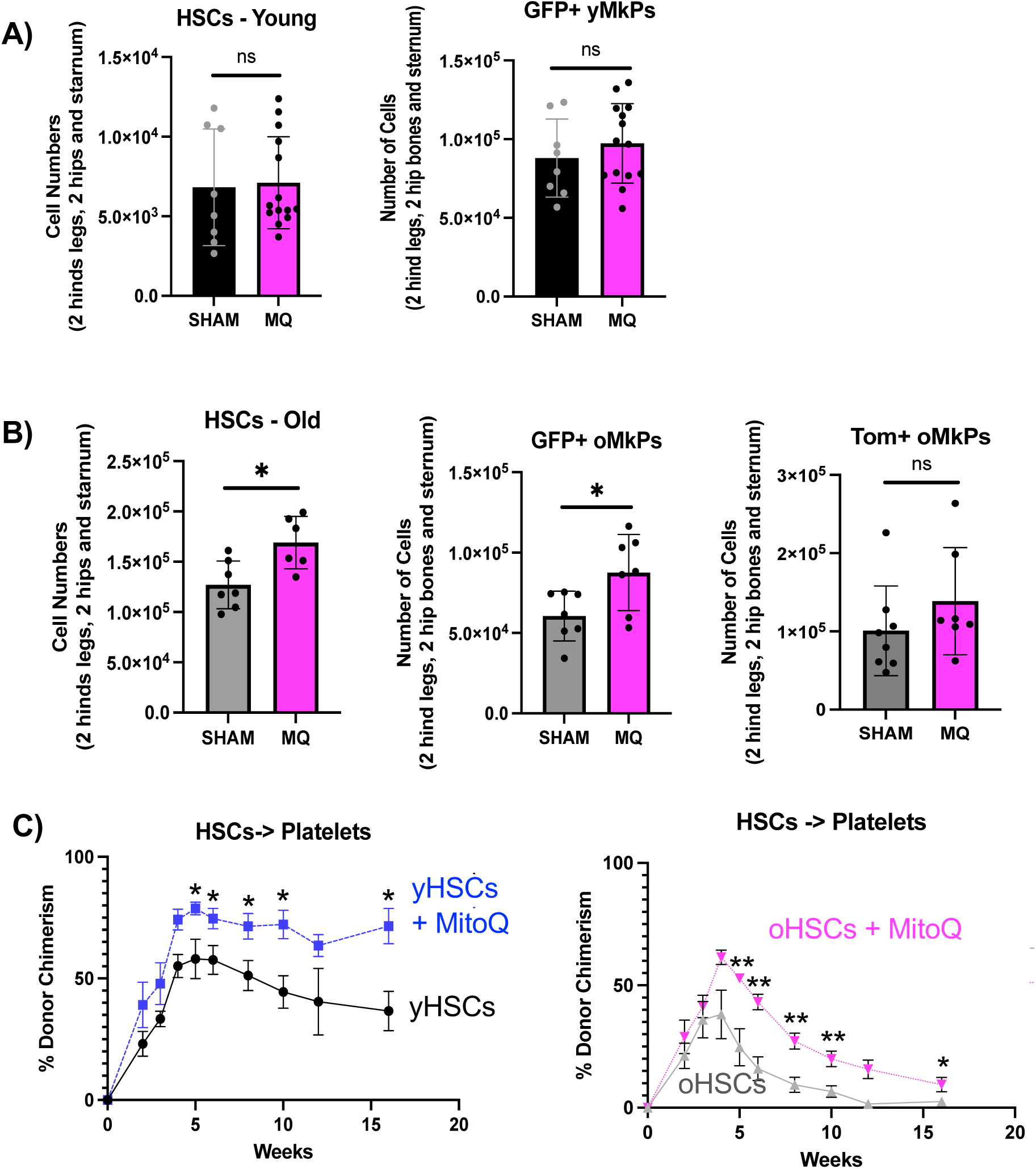
Changes in HSC and MkP numbers in old, but not young, mice upon MitoQ treatment. **A.** HSC and MkP numbers did not change upon MitoQ treatment of young mice. Changes in the number of HSCs and GFP+ MkP in young upon MitoQ treatment. Student’s t-test. (n = 8-12 in 3 independent experiments). **B.** oHSC and GFP+, but not Tom+, oMkPs increase upon MitoQ treatment of old mice. Changes in the number of HSCs, GFP+ MkPs and Tom+ MkPs upon MitoQ treatment in old mice. Student’s t-test. (n = 6-7 in 3 independent experiments; *P < 0.05). **C.** Enhanced Plt reconstitution by both young and old MitoQ-treated HSCs. %Donor chimerism upon transplantation of HSCs isolated from young and old control and MitoQ-treated mice. Donor-derived Plts in PB of recipients for 16 weeks post-transplantation presented as percent donor chimerism. Data represent mean ± SEM of 3 independent experiments, Statistics: unpaired two-tailed t-test. *P<0.05, **P<0.005, ***P<0.0005.

Thus, to assess the functional capacity of HSCs upon MitoQ treatment, we executed transplantation assays, the gold standard for evaluating the functional output of HSCs and progenitors. 200 HSCs isolated from both young and old MitoQ-treated or untreated mice were transplanted into sublethally irradiated young mice. PB reconstitution was monitored until the 16-week endpoint, when BM was also analyzed for donor cell engraftment. Since recipient mice were never exposed to MitoQ, any changes in chimerism would result from intrinsic changes in old or young populations from donors that were treated in vivo prior to cell isolation and transplantation. Interestingly, transplantation of HSCs isolated from young MitoQ-treated mice demonstrated a marked improvement in platelet reconstitution compared with HSCs from young sham-treated controls. This is reminiscent of the increased in situ Plt production in young MitoQ-treated mice (**Fig 4C**). Because Plt production in old MitoQ-treated mice decreased (**Fig 4C-D**), we were surprised that HSCs isolated from old MitoQ-treated mice displayed a significant increase in Plt production upon transplantation (**Fig 5C**). Comparison across age groups revealed that young HSCs isolated from sham treated mice showed robust reconstitution of Plts compared to sham-treated oHSCs, as expected^3,4^ (Fig 5C). This underscores the well-established decline in regenerative potential that accompanies HSC aging. While MitoQ treatment led to improved Plt output of both young and old HSCs, MitoQ-treated yHSCs maintained more robust and maintained Plt reconstitution compared to MitoQ-treated oHSCs. This suggests that MitoQ can act beyond restoration of oHSC to yHSC capacity, revealing an important role of mitochondrial fitness in HSC reconstitution potential, irrespective of age.

## DISCUSSION

### Aging-specific MkPs have superior mitochondrial functionality

Aging profoundly impacts blood cell production, especially Plt generation, leading to both qualitative and quantitative changes in Plt production^3,4^. This study focused on changes in mitochondrial function in aged MkPs. We demonstrated that the emergence of the aging-progressive bypass pathway to Plt generation upon aging is associated with differential mitochondrial regulation. Under young, steady-state conditions, this pathway remains largely dormant, while aging leads to the accumulation of age-specific MkPs (**Fig 1**). With aging, deterioration of mitochondrial structure, function, and dynamics has become increasingly evident for various tissues. This includes impaired mitochondrial energy production, which contributes to the accumulation of dysfunctional mitochondria. Although several studies have associated decreased energy production and elevated ROS with oHSCs, it is not clear whether these mitochondrial changes are causal drivers of the age-related increase in Plt production.

Surprisingly, while oHSCs showed a reduction in mitochondrial capacity upon aging, MkP MMP and ATP levels increased with age. This was primarily due to the emergence of ncMkPs. We observed that age-specific ncMkPs have superior mitochondrial functionality compared to both young and co-existing old cMkPs, including elevated MMP and ATP levels and lower ROS (**Fig 2 and 3**). These findings support the notion that reduced MMP is linked to cellular deterioration^9,34^, whereas higher MMP may underlie the superior repopulation and expansion capacity of ncMkPs compared to co-existing aged cMkPs ^4,15^.

### Mitochondrial enhancement differentially alters platelet counts in young and old mice

Several studies have provided evidence that targeted modulation of mitochondrial function can reverse the key hallmarks of oHSCs^9,18,25,35^, suggesting that restoring mitochondrial homeostasis has therapeutic potential. Evidence suggested that these effects are mediated by direct alteration of MMP and mitochondrial biology at the cellular level. Surprisingly, enhancement of mitochondrial activity elicited opposing effects on Plt counts in Y versus O mice. In young mice, MitoQ treatment increased Plt production, whereas Plt counts decreased in old mice; this reduction was selective for Tom+ oPlts (**Fig 4**). In contrast, alterations in erythroid, B, and myeloid cells were consistent across young and old mice (**Fig 4**). These findings show that enhancing mitochondrial function differentially affects different lineages. The short MitoQ treatments period versus the relatively long half-lives of erythroid and B cells make it unlikely that their increased and decreased numbers, respectively, are a reflection of HSC fate choice. Instead, our data suggest that more mature erythroid and B lineage cells respond to mitochondrial enhancement. Plt counts may similarly be regulated by MitoQ-responsive progenitors rather than by HSCs, but the short Plt half-life of ∼5 days and rapid Plt replacement kinetics in vivo^4^ makes it possible that HSCs contribute to the observed alterations in the Plt pool. Interestingly, we observed an increase in oHSCs, but not yHSCs, upon MitoQ treatment, along with an unexpected increase in old cMkPs, but not ncMkPs (**Fig 5**). The differential responses of young and old HSCs and MkPs to mitochondrial enhancement underscore that the balance in mitochondrial adaptation is central to maintaining hematopoietic homeostasis.

Perturbing this balance, even transiently, can alter lineage composition, suggesting that targeted modulation of mitochondrial function may represent a powerful but nuanced strategy to correct age-related hematopoietic dysfunctions. While more research is needed to pinpoint the responsive cell type(s), MitoQ may act to restore redox balance and reduce mitochondrial stress, effectively attenuating the overproduction of Plts upon aging.

### Enhancement of mitochondrial capacity increases platelet repopulation by both young and old HSCs

Mitochondrial enhancement has been proposed to act as a rejuvenating agent that can restore aged cells to a more youth-like behavior. Our results support the notion that HSC functionality, measured by reconstitution capacity upon transplantation (**Fig 5**), is increased by MitoQ. Intriguingly, we found that the repopulation activity of yHSCs was also increased. This suggests that mitochondrial enhancement of HSC engraftment is aging-independent and can act beyond restoration to a youthful baseline. These short-term MitoQ effects on HSCs are at least partially cell intrinsic and, intriguingly, can be maintained upon stem cell transplantation with long-term consequences for function. While the gain in repopulation potential was age-independent, mitochondrial regulation of megakaryopoiesis in situ was age-dependent. Thus, while mitochondrial enhancement may confer broad rejuvenative effects on HSC repopulation capacity, it may also suppress selectively amplified outputs, such as thrombopoiesis, that arises upon aging. Mitochondrial therapeutics may therefore be capable of mitigating age-related hematopoietic decline and its downstream consequences, including thrombotic risk.

## MATERIALS AND METHODS

### Mice

All animals were maintained under approved IACUC guidelines in the AAALAC accredited vivarium at University of California Santa Cruz. The following mice were utilized for these experiments: C57Bl6 (JAX, cat# 664), aged C57Bl6 (NIH-ROS), and male FlkSwitch mice (Flt3-Cre x mTmG mice). Young mice were between 8-16 weeks of age and old mice were 20+ months of age, except old transplant recipient mice which were 18+ months of age. All WT C57Bl6 mice were randomized based on sex.

### Flow Cytometry

Bone marrow stem and progenitor cell populations and mature cell subsets were prepared and stained as previously described^3,4,15^. Briefly, the long bones (femur and tibia) from mice were isolated, crushed with a mortar and pestle, filtered through a 70μm nylon filter and pelleted by centrifugation to obtain a single cell suspension. Cell labeling was performed on ice in 1X PBS and 2% FBS. HSCs (Lin-cKit+Sca1+Flk2-Cd150+Tom+ or Lin-cKit+Sca1+Flk2-Cd150+CD48-) or MkPs (Lin-cKit+Sca1-Cd41+Cd150+Tom+ or GFP+ or Lin-cKit+Sca1-CD41+CD150+CD48-/+CD321-/+) from young or old FlkSwitch and WT mice were analyzed from unfractionated samples or isolated from c-Kit-enriched BM with CD117-microbeads (Miltenyi) using a FACS ARIA II (Becton Dickinson, San Jose, CA) as previously described. Cells with no history of Cre expression were defined as Tom+GFP-(“unfloxed”) cells, whereas cells with current or a history of Cre expression were defined solely by GFP expression, based on our previous demonstration that Tom+GFP+ cells have excised the loxP-flanked Tomato cassette; that Tom+GFP+ and Tom-GFP+ cells are functionally indistinguishable; and well-accepted field standards of loxP-stop-loxP-inducible single-color reporters.

### Nucleic acid isolation

The mtDNA and cellular DNA were isolated from sorted cells with DNeasy Blood and Tissue Kit (#69504 Qiagen) according to the manufacturer’s protocol. Briefly, sorted cells were resuspended in 200 μl PBS to which 20 μl proteinase K and RNaseA were then added. After the addition of 200 μl buffer AL, the samples were vortexed and kept at 56C for 10 mins. Ethanol (100%, 200 μl) was then added and the samples vortexed again. The mixture was pipetted into a spin column which was then centrifuged at 6000×g for 1 min on top of a collecting tube. The column was then rinsed twice, first with 500 μl buffer AW1 (centrifugation at 6000×g for 1 min) and second with 500 μl buffer AW2 (centrifugation at 20,000×g for 3 min). Nucleic acids were eluted with 200 μl buffer AE (centrifugation at 6000×g for 1 min). All centrifugations were performed at room temperature. Nucleic acid concentrations and purity were assessed with a NanoDrop spectrophotometer.

### qPCR (mtDNA & ncDNA)

Real-time quantitative PCR (qPCR) to quantify mtDNA and ncDNA was performed on 20-30 ng nucleic acids, directly after the nucleic acid isolation step, by using the SYBR Green No ROX Kit (#QT650-Bioline) and with the following program in Quant Studio 6 Flex (Thermo): 2 min at 50C, 5 min at 95C, 45 cycles of 15s at 95C, 15s at 59C and 15s at 72C. ncDNA was quantified by using the following primers: Hexokinase 2 (HK2) Forward Primer: 5’-GCCAGCCTCTCCTGATTTTAGTGT-3’, HK2 Reverse Primer: 5’-GGGAACACAAAAGACCTCTTCTGG-3’. Total mtDNA was quantified by amplifying a DNA domain within the D-loop of mtDNA by using the follow-ing primers: 16S rRNA Forward Primer: 5’-CCGCAAGGGAAAGATGAAAGAC-3’, 16S rRNA Reverse Primer: 5’-TCGTTTGGTTTCGGGGTTTC-3’, NADH Dehydrogenase subunit I (ND1) Forward Primer: 5’-CTAGCAGAAACAAACCGGGC-3’, ND1 Reverse Primer: 5’-CCGGCTGCGTATTCTACGTT-3’. Analysis has been done according to this published article^36^.

### TMRE-Mitochondrial Membrane Potential Assay and Mitotracker Green (MTG)

To assess mitochondrial mass and MMP, cells were subjected to sequential cell surface and mitochondrial staining. Following completion of surface marker labeling, cells were washed once in cold SM (PBS + 2% FBS) and subsequently incubated at 37°C for 30 minutes in PBS containing one of the following mitochondrial probes: MTG (50 nM; Thermo Fisher Scientific) or TMRE (100 nM; Thermo Fisher Scientific). Staining was performed in the presence of verapamil (50 µM; Sigma-Aldrich, #V4629), which was prepared as a 10 mg/mL stock solution in DMSO and diluted in PBS immediately before use. After staining, cells were washed twice with SM (400 × g, 5 minutes, 4°C) and resuspended in fresh buffer for flow cytometric acquisition. Data were collected using a BD FACS AriaII or Ctyoflex and analyzed using FlowJo (BD Biosciences). To confirm mitochondrial specificity and validate the sensitivity of each probe, cells were treated with the mitochondrial uncoupler carbonyl cyanide m-chlorophenyl hydrazone (CCCP; 50 µM) for 30 minutes following MMP or MTG staining.

### ROS and mitochondrial superoxide

Intracellular ROS levels were quantified by flow cytometry following cell surface staining using the DCFDA/H₂DCFDA Cellular ROS Assay Kit (Abcam, Cat# ab113851), according to the manufacturer’s instructions. Briefly, cells were resuspended in 1X assay buffer containing 20 µM DCFDA and incubated for 30 minutes at 37C protected from light. After incubation, cells were washed once with assay buffer and immediately analyzed by flow cytometry to prevent oxidation artifacts or dye leakage. As previously reported, VP can reduce DCFDA fluorescence across bone marrow lineages by inhibiting dye uptake; therefore, VP was excluded from all DCFDA assays. To verify dye responsiveness and signal fidelity, cells were exposed to tert-butyl hydroperoxide (TBHP), which robustly increased DCFDA fluorescence, confirming that the chosen concentration and incubation time effectively captured changes in intracellular ROS. Mitochondrial-specific superoxide production was quantified using the MitoSOX™ Red Mitochondrial Superoxide Indicator (Thermo Fisher Scientific, Cat# M36008). Cells were incubated in PBS containing 5 µM MitoSOX Red for 30 minutes at 37C in the presence of 50 µM verapamil (Sigma-Aldrich, Cat# V4629), prepared in PBS from a 10 mg/mL DMSO stock solution, to minimize dye efflux via multidrug resistance transporters. Following staining, cells were washed once (400 × g, 5 minutes) and resuspended in SM for immediate flow cytometric acquisition.To confirm assay sensitivity and specificity, cells were treated with TBHP, which induced a pronounced increase in MitoSOX fluorescence, validating the probe’s responsiveness to mitochondrial superoxide generation under oxidative stress conditions.

### ATP content

Intracellular adenosine triphosphate (ATP) levels were quantified using the Luminescent ATP Detection Assay Kit (Abcam, Cat# ab113849) following the manufacturer’s protocol with minor modifications to optimize for low cell numbers. Briefly, 1,500 HSCs, MkPs, or subpopulations of MkP were FACS-sorted directly into individual wells of a 96-well white, opaque flat-bottom plate optimized for luminescence detection (Costar, Cat# 3615) containing 100 µL SM. Following sorting, cells were gently washed once with PBS to remove residual serum proteins that could interfere with the enzymatic reaction. Next, 50 µL of detergent solution was added to each well, and plates were incubated for 5 minutes on an orbital shaker (600–700 rpm) at room temperature, protected from light, to ensure complete cell lysis and ATP stabilization.

Subsequently, 50 µL of the luciferase substrate solution was added to each well, followed by another 5-minute incubation under identical conditions. To ensure stable luminescent signal detection, plates were dark-adapted for 10 minutes by covering them with aluminium foil prior to reading. Background readings from blank wells containing only buffer and reagents were subtracted from all measurements. All samples were analyzed in a luminescence plate reader in biological triplicates and technical duplicates to ensure consistency.

### MitoQ experiments

Mitoquinol was administered through intra-peritoneal injection (2 mg/kg/d <u>body weight</u>, <u>Cayman</u> Cat# 89950) for 5 days in either young or old mice. Control mice were injected with vehicle(DMSO) dissolved in HBSS. Mice were bled on days 0, 3, and 5 to assess changes in platelet and mature blood cell lineages by flow cytometry. Weights were taken daily of both treated and non-treated animals. Both control and treated animals were euthanized by CO2 inhalation on Day 5 post final injection of MitoQ. Cell populations analyzed in the peripheral blood and bone marrow.

### Transplantation

For the sort, briefly, the long bones (femur and tibia), hips and sternum from control and the treated mice were isolated separately, crushed with a mortar and pestle, filtered through a 70μm nylon filter and pelleted by centrifugation to obtain a single cell suspension. Cell labeling was performed on ice in 1X PBS with 5 mM EDTA and 2% FBS. HSC (Lin-/cKit+/Sca1/CD150+/Flk2-/Tom+) and MkP (Lin-/cKit+/Sca1-/CD41+/Slam+), separated into cMkPs and ncMkPs using Tom/GFP or CD48/CD321, as described, were double sorted from FlkSwitch mice post ckit - enrichment. 200 HSC or 22,000 MkP were transplanted into sub-lethally irradiated mice (5Gy) with a Multi-Rad CP-160. Beginning two weeks post-transplantation, recipient mice were bled weekly via the tail vein. Peripheral blood was analyzed for donor chimerism as described above and in previous studies.

## ACKNOWLEDGEMENTS

We thank Dr. M.G.E. Rommel for critical review of the manuscript, the University of California Santa Cruz Institute for the Biology of Stem Cells (IBSC) flow cytometry (RRID:SCR_021149), stem cell culture (RRID:SCR_021353), and vivarium facilities. This study was supported by the National Institutes of Health (NIH) National Institute on Aging (NIA) R01AG062879 to E.C.F., NIH Eunice Kennedy Shriver National Institute of Child Health and Human Development (NICHD) T32HD108079 and California Institute for Regenerative Medicine (CIRM) EDUC4-12759 to S.C., by NIH S10OD030423 to ECF/UCSC and by CIRM Shared Stem Cell Facilities (CL1-00506), CIRM Major Facilities (FA1-00617-1) awards to UCSC.

## AUTHOR CONTRIBUTIONS

S.C. conceived of the study. S.C and E.C.F. performed data analysis and wrote the manuscript. S.C., S.S.B. and S.H. performed all experiments.

## DECLARATION OF INTERESTS

The authors declare no competing interests.

## Notes

### Competing Interest Statement

The authors have declared no competing interest.

